# Molecular arms race in WHO elements, a category of homing genetic elements distinct from inteins and introns

**DOI:** 10.64898/2026.08.13.744650

**Authors:** Matthieu Osborne, Ludovic Monnin, Kenneth H. Wolfe

**Affiliations:** Conway Institute and School of Medicine, University College Dublin, Dublin 4, Ireland

**Keywords:** homing endonucleases, LAGLIDADG endonucleases, aldolase, genetic conflict, yeast, *Torulaspora*

## Abstract

Homing genetic elements are selfish elements that insert themselves into a specific site in a host gene without disrupting its function. They spread through the population because the element codes for an endonuclease that cleaves alleles of the host gene that do not contain the element, leading to DNA repair by gene conversion that increases the element’s frequency. Most homing genetic elements in eukaryotes are either self-splicing introns or inteins but we recently discovered a third category, called WHO elements, in the budding yeast genus *Torulaspora*. WHO elements code for endonuclease proteins with LAGLIDADG motifs and a zinc finger domain, and are related to the mating-type switching endonuclease HO. Their host gene is the aldolase gene *FBA1*, which is essential. Clusters of up to 9 diverse *WHO* endonuclease genes are found downstream of *FBA1* in different isolates of *Torulaspora*. Here, we show that there is a genetic conflict between WHO endonucleases and their target site in *FBA1.* Different alleles of *FBA1* vary in their sensitivity or resistance to cleavage by individual WHO endonucleases. We show that a WHO endonuclease recognizes a 28-bp sequence in *FBA1* and does not tolerate much sequence variation, but also that this region of *FBA1* has experienced positive selection for sequence diversification to evade cleavage. WHO endonucleases and their target site in *FBA1* are therefore engaged in an arms race in which each WHO element is under selection to home into other elements, while avoiding being homed into.

**Significance Statement:** WHO elements are a recently discovered type of homing genetic element in yeasts, targeting the aldolase gene *FBA1*. Rather than disrupting *FBA1* when they integrate, WHO elements instead replace the 3’ half of the gene with an alternative *FBA1* 3’ half. Each WHO element consists of an endonuclease gene and a version of the 3’ half of *FBA1*, and there is high sequence diversity in both genes. We show that there is an evolutionary arms race between WHO endonucleases and their target site in *FBA1*, which has resulted in rapid evolution of both genes and the formation of clusters of *WHO* elements at the *FBA1* locus.

## Introduction

Homing genetic elements are selfish elements capable of inserting themselves into a specific target site in the genome (1, 2). Most homing elements are found in prokaryotes or bacteriophages but there are some in eukaryotes, including the omega self-splicing intron which targets the 21S rRNA gene of the *Saccharomyces cerevisiae* mitochondrial genome, and the VDE intein which targets the *S. cerevisiae* nuclear gene *VMA1* (3, 4). These elements occupy a site within their host gene but they do not interfere with its expression because they are removed during transcription (for introns) or post-translationally (for inteins). The host gene is usually polymorphic, so that alleles containing the homing element, and alleles lacking it, are both present in the population. We refer to these as occupied and unoccupied alleles.

Homing elements encode endonucleases that can recognize and cleave a target DNA site in unoccupied alleles of their host gene. The endonucleases encoded by known homing elements in the nuclear genomes of eukaryotes are members of the LAGLIDADG family (2, 5, 6). Their target site is typically more than 18 bp long and occurs only once in the host’s genome. In a non-haploid context, the dsDNA break in the cleaved allele can be repaired by using the other allele, i.e. the occupied allele, as a template. This process results in gene conversion and increases the homing element’s frequency in the population. The endonuclease produced by an occupied allele can cleave unoccupied alleles, but it cannot cleave its own allele because the element disrupts the target site when it is inserted, making the occupied allele resistant to self-cleavage.

When our laboratory sequenced the genome of the yeast *Torulaspora delbrueckii* (strain CBS1146) in 2011 (7), we found an unusual cluster of LAGLIDADG family genes and pseudogenes immediately downstream of the gene *FBA1*, which codes for fructose-1,6-biphosphate aldolase, a glycolysis enzyme. As well as having an endonuclease domain with two LAGLIDADG motifs, the proteins encoded by these genes also have a zinc finger domain at the C-terminus. This domain organization is unusual and is not seen in any other LAGLIDADG endonucleases except for HO, the endonuclease that cleaves the yeast *MAT* locus during mating-type switching (8), so we named them *WHO* genes (for “<u>w</u>eird *<u>HO</u>*-like” genes) (9). The cluster in *T. delbrueckii* CBS1146 contains three intact *WHO* genes and three *WHO* pseudogenes (Fig. 1). Although the *WHO* genes have a LAGLIDADG domain typical of homing element endonucleases, they do not seem to interrupt a host gene so they are evidently not part of an intein or an intron-encoded homing element (9). The *WHO* genes are also diverse in sequence, with only 55% amino acid sequence identity between the most similar pair in the cluster of *WHO* genes in *T. delbrueckii* CBS1146. We found that there was extensive structural variation among the *WHO* gene clusters present in different isolates of *T. delbrueckii*, with clusters containing between 2 and 9 different *WHO* genes or pseudogenes, and that the clusters also vary extensively among species in the genus *Torulaspora* (Fig. 1; (9, 10)).

**Figure 1.**
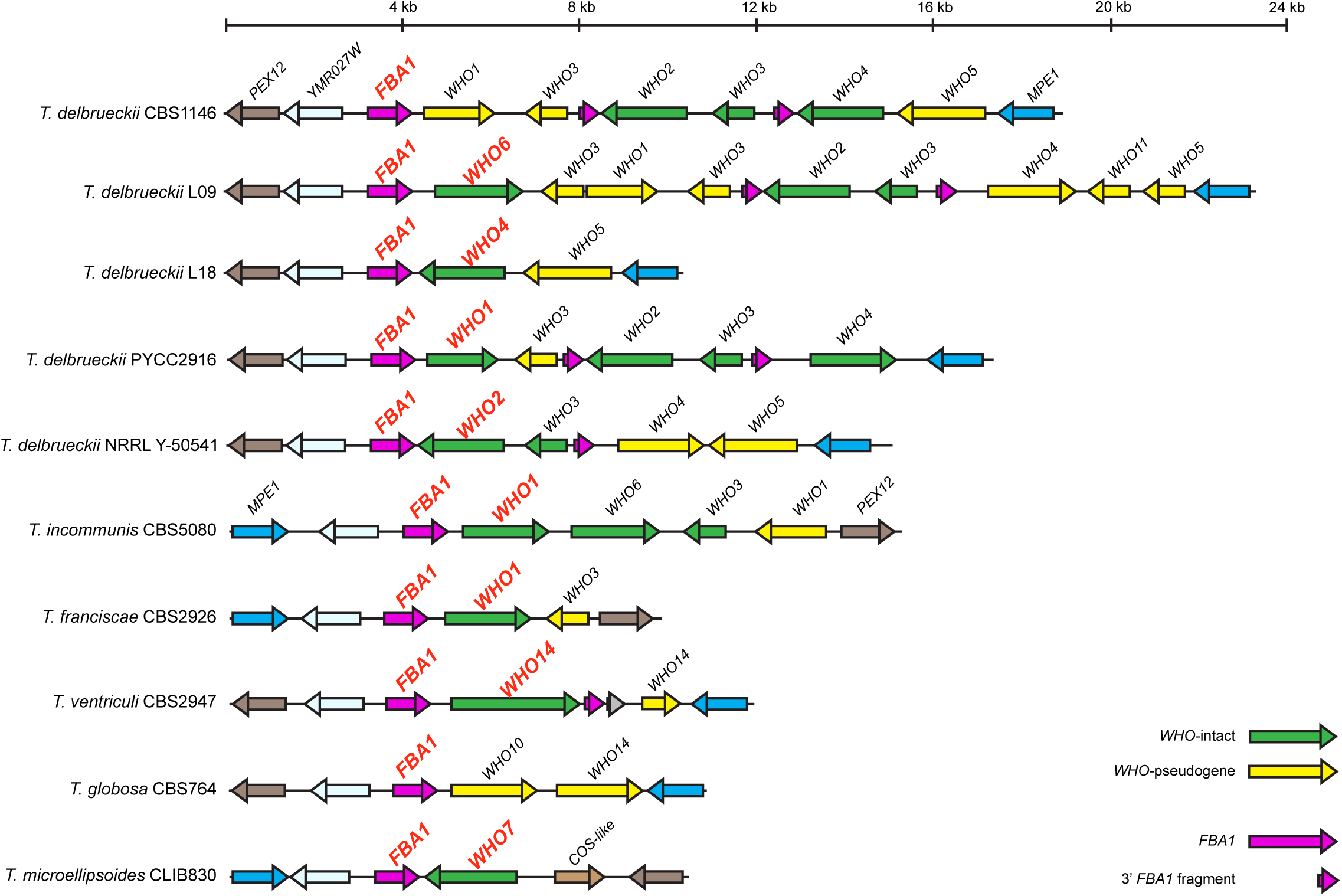
Structure of *WHO* gene clusters in *Torulaspora* strains and species. The *WHO* genes and *FBA1* alleles used in this study are named in red. Green arrows show intact *WHO* genes, and yellow arrows show *WHO* pseudogenes. *FBA1* full-length genes and 3’ fragments are shown in magenta. In *T. delbrueckii* CBS1146 and *T. globosa* CBS764 the *WHO* gene immediately downstream of *FBA1* is a pseudogene; we investigated *FBA1* alleles, but not *WHO* genes, from these strains. Species are named according to the classification proposed by Silva *et al*. (10).

By experimental analysis, we discovered that the target site for cleavage by WHO endonucleases is located in *FBA1*. We found that a WHO endonuclease from one isolate of *T. delbrueckii* (WHO6, from strain L09) was able to cleave the *FBA1* allele from a different isolate (CBS1146), but it could not cleave the *FBA1* allele from strain L09 itself (9). The cleavage site is near the center of the gene and divides *FBA1* approximately into two halves. The clusters of *WHO* genes and pseudogenes found downstream of the full-length *FBA1* gene in *Torulaspora* often contain additional copies of the 3’ half of *FBA1*, beginning at the cleavage site (Fig. 1). When comparing the sequences of *FBA1* alleles among *T. delbrueckii* isolates, we also found that the sequence diversity in the 3’ half of the gene is much greater than in the 5’ half, and that the two halves of the gene have different phylogenies (9).

These observations led us to hypothesize that *WHO* genes form part of a new type of homing genetic element, which we named WHO elements. The proposed organizational structure of WHO elements is quite different from the structure of inteins and homing introns, even though all three elements share the same molecular mechanism of homing (9, 11). Under this hypothesis, *FBA1* is the host gene of WHO elements (Fig. 2A). A complete WHO element consists of a *WHO* endonuclease gene and the 3’ half of the full-length, functional, *FBA1* gene that lies immediately upstream of it (Fig. 2B). We hypothesized that each WHO endonuclease is unable to cleave the *FBA1* sequence that lies directly upstream of its own gene in the same WHO element (so the element does not cleave itself), but it may be able to cleave other alleles of *FBA1* that have different sequences in their 3’ half. We hypothesized that homing occurs in diploid *Torulaspora* cells that are heterozygotes for the presence/absence of a WHO element. During homing, the WHO endonuclease encoded by one allele cleaves the *FBA1* gene located on the other allele, leading to DNA repair, overwriting of the unoccupied allele by the occupied allele, and formation of a chimeric *FBA1* gene with a new 3’ half (Fig. 2B). Homing could also occur if a diploid is heterozygous for two different WHO elements (Fig. 2C). We hypothesized that the clusters of WHO elements downstream of *FBA1* have been formed by successive homing by different elements with different endonuclease specificities and different 3’-*FBA1* sequences (Fig. 2C,D). Within each cluster, the *WHO* gene located closest to the full-length *FBA1* gene is expected to be the one that was inserted most recently (9, 11). The *WHO* genes located further away, and the extra copies of the 3’ half of *FBA1*, are relics from older homing events and are liable to decay into pseudogenes or to become deleted, similar to the evolutionary decay that occurs in other homing elements (12, 13). We proposed an ‘arms race’ model for how WHO endonucleases and their target site in *FBA1* coevolve (11). Arms races between two interacting genes, resulting in rapid evolution of both of them, are a frequent outcome of genetic conflicts (1, 14, 15).

**Figure 2.**
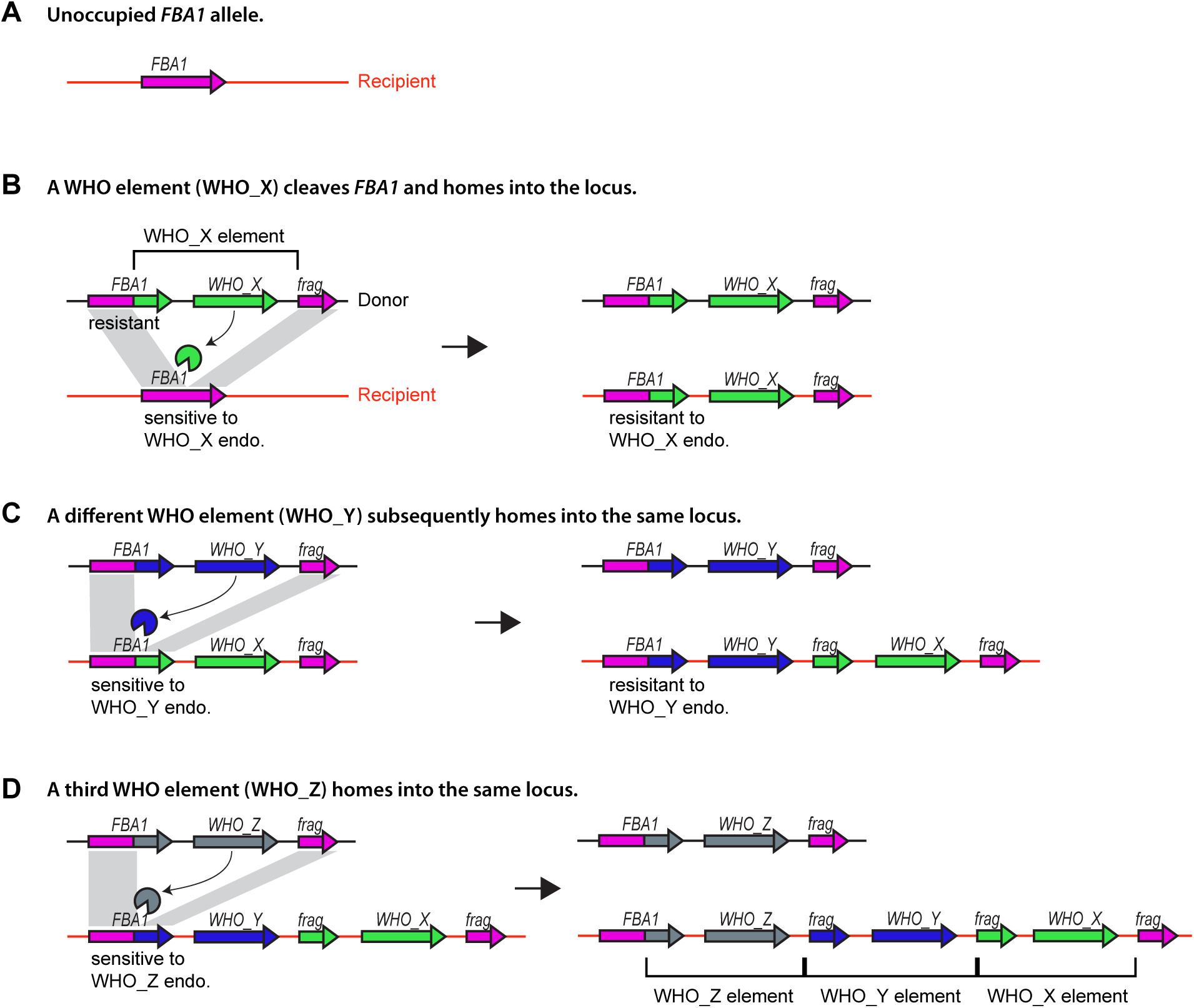
Hypothesis for WHO element structure, homing, and cluster formation. Sequential homing of three hypothetical WHO elements (WHO_X, WHO_Y and WHO_Z) into the same locus on a recipient chromosome (red line) is depicted. **(A)** The locus is initially unoccupied and contains a single *FBA1* gene. **(B)** In a heterozygous diploid formed by mating in the population, the endonuclease encoded by a WHO_X element on the donor chromosome cleaves the *FBA1* gene of the recipient chromosome and homing occurs. Homing replaces the 3’ half of the full-length *FBA1* gene on the recipient chromosome, which makes it resistant to cleavage by WHO_X endonuclease but may leave it sensitive to other WHO endonucleases. The 3’ part of the original *FBA1* gene on the recipient chromosome is left as a fragment downstream of the integrated WHO element. **(C)** Generations later, a diploid heterozygous for WHO_X and WHO_Y elements is formed by mating in the population. The *FBA1* allele on the recipient chromosome is sensitive to cleavage by the WHO_Y endonuclease, so the WHO_Y element homes into it, upstream of the WHO_X element. For simplicity, we assume here that WHO_X endonuclease cannot cleave the *FBA1* gene of the donor chromosome. **(D)** Later, a WHO_Z element homes into the same locus, enlarging the cluster. At all stages, the WHO element immediately downstream of the full-length *FBA1* gene is the one that integrated most recently. The ones further away are older and often acquire mutations that turn their *WHO* genes into pseudogenes or delete the *FBA1* fragments.

In this paper, we test some of the key predictions of this hypothesis. We report that WHO endonucleases have varying abilities to cleave different alleles of *FBA1*. We find that the DNA sequence in *FBA1* recognized by a WHO endonuclease is of similar length (28 bp) to the recognition sequences of the related LAGLIDADG endonucleases HO and VDE, but is less degenerate. We find that the recognition site in *FBA1* is evolving quickly and has been subject to positive selection. Each of these results is consistent with the hypothesis that an arms race is underway between WHO endonucleases and their target site in *FBA1*, due to the opposing selective pressures acting on these two neighboring genes that together form a WHO element.

## Results

### Assay system based on β-estradiol induction

To investigate the interactions between WHO endonucleases and *FBA1* target sites, we assayed the ability of 8 different WHO endonucleases to cleave 10 different alleles of *FBA1*. We used a system in which the expression of a heterologous gene in *S. cerevisiae* can be induced by addition of the mammalian hormone β-estradiol to the media, via a synthetic transcription factor that drives a synthetic promoter (16, 17). We first integrated the synthetic transcription factor gene at the *his3* locus of the haploid *S. cerevisiae* strain BY4742 (Fig. 3A). We then integrated each *Torulaspora WHO* gene under the synthetic promoter at the *leu2* locus (Fig. 3B), to make 8 WHO-expressing strains. Finally, we used CRISPR-Cas9 at the *ho* locus (Fig. 3C) to integrate an 83-bp target site region, centered on the site known to be cleaved by WHO6, from each of the 10 *Torulaspora FBA1* sequence variants into each of the 8 WHO-expressing strains, to make 80 test strains. When induced by β-estradiol, the WHO protein can cleave the *FBA1* target site, creating a double strand break (DSB) (Fig. 3D). If a DSB is created, most cells will die because accurate DNA repair leads to a futile cycle of re-cleavage and results in a growth defect, at least until a spontaneous resistance mutation arises in the culture (9, 18).

**Figure 3.**
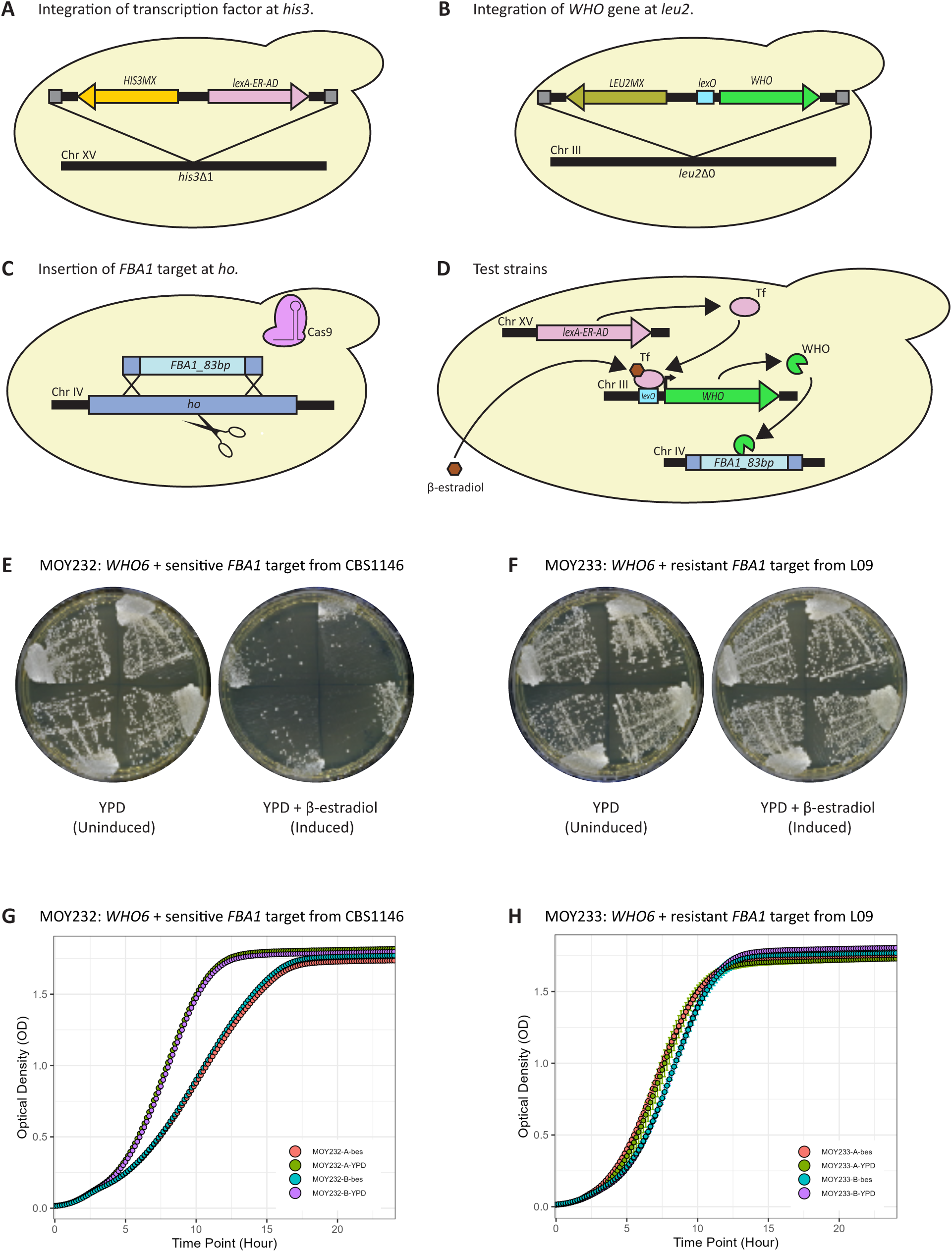
Beta-estradiol induction system used to assess WHO endonuclease target specificity. **(A-D)** Construction of the system. **(A)** The synthetic β-estradiol activated transcription factor gene *LexA-ER-AD* is integrated at the *his3* locus of haploid *S. cerevisiae* strain BY4742. **(B)** A *WHO* endonuclease gene with a *lexO* promoter controlled by the synthetic transcription factor is integrated at the *leu2* locus. **(C)** The sequence of an 83-bp region around the target site in a *Torulaspora FBA1* gene is integrated at the *ho* locus by CRISPR-Cas9 editing. **(D)** When β-estradiol is present in the media, it enters the cell and activates the transcription factor (Tf), enabling it to enter the nucleus and bind to *lexO* which turns on transcription of the *WHO* gene. If the WHO endonuclease successfully cleaves the 83-bp *FBA1* target, the strain’s growth rate will be reduced. **(E-H)** Verification of the system by using WHO6 endonuclease and known sensitive and resistant target sequences, in solid and liquid media assays. *S. cerevisiae* strain MOY232 contains the *WHO6* gene and the sensitive *FBA1* target sequence from *T. delbrueckii* CBS1146, while MOY233 contains the *WHO6* gene and the resistant *FBA1* target sequence from *T. delbrueckii* L09 (9). When induced on β-estradiol, MOY232 shows a growth defect compared to MOY233, both on solid media (quadrants show 4 biological replicates) and in liquid growth assays (2 biological replicates are shown).

We validated the system by verifying that WHO6 endonuclease can cleave the *FBA1* allele from *T. delbrueckii* CBS1146 but not from *T. delbrueckii* L09, as was shown previously by using a different expression system (9). When streaked to YPD agar plates containing β-estradiol, *WHO6*-expressing strains die if they contain the CBS1146 *FBA1* allele, but not if they contain the L09 *FBA1* allele (Fig. 3E,F). To obtain a quantitative readout of growth reduction, we used liquid cultures and calculated the difference (Δ*k*) between the slopes of the growth curves of uninduced and induced cultures, at their maximal growth rates (Fig. 3G,H). When *WHO6* was induced in a strain containing the sensitive CBS1146 *FBA1* allele, the difference in slopes was Δ*k* = 0.36, whereas there was no reduction of growth rate in a strain containing the resistant L09 *FBA1* allele (Δ*k* = 0.00).

### Allele-specific cleavage of *FBA1* by WHO endonucleases

Our hypothesis for the structure and homing mechanism of WHO elements is only valid if cleavage of *FBA1* by WHO endonucleases shows allelic variation, i.e. if each WHO endonuclease can cleave some *FBA1* alleles but not others, and if each *FBA1* allele is susceptible to cleavage by some WHO endonucleases but not by others. Moreover, each *FBA1* allele should be resistant to cleavage by the product of the *WHO* gene located directly beside it.

We tested this hypothesis by using eight WHO endonuclease genes, from five species of *Torulaspora* including four isolates of *T. delbrueckii* (Fig. 1; Fig. S1). The endonucleases encoded by these genes have 33% to 83% amino acid sequence identity and were chosen to get a broad perspective on WHO protein specificity. There are approximately 14 phylogenetic families of WHO endonuclease genes, including some from species other than *Torulaspora* and some that were only found as pseudogenes (9). The sequences we used include representatives of six families: WHO1, WHO2, WHO4, WHO6, WHO7, and WHO14. We included three *WHO1* genes that are quite similar (76% to 83% amino acid identity) but which come from different species: *T. delbrueckii*, *T. franciscae*, and *T. incommunis* (Fig. S1). Each of the *WHO* genes we assayed is located beside a full-length *FBA1* gene, and they are all located in *WHO* gene clusters except in *T. microellipsoides* which has only one *WHO* gene (Fig. 1).

We assayed the sensitivity of 10 alleles of *FBA1* to each of the eight WHO endonucleases. They include the full-length *FBA1* genes found beside the eight *WHO* genes that we tested (Fig. 1), as well as the *FBA1* genes from *T. delbrueckii* strain CBS1146 and *T. globosa* CBS764. The latter two *FBA1* genes are located beside *WHO* pseudogenes (*ψWHO1* and *ψWHO10* respectively; Fig. 1), which we assumed to be incapable of coding for active endonucleases. For each *FBA1* allele, we tested the ability of WHO endonucleases to cleave an 83-bp sequence centered on the known WHO6 cleavage site at nucleotide positions 666-669 of *FBA1* (9), using quantification of growth reduction as a proxy for cleavage. Nucleotide sequence identity among the 10 *FBA1* alleles ranges from 80% to 98% across the whole gene, and from 65% to 96% in the 83-bp target site region tested.

The resulting matrix of pairwise interactions between *WHO* genes and *FBA1* alleles is shown in Figure 4A. Some pairs show growth reduction almost as strong as was seen between WHO6 and the *FBA1* allele from *T. delbrueckii* CBS1146 (Fig. 3), whereas other pairs show little or no reduction. We chose a cutoff value of Δ*k* ≥ 0.15 as an indication of a significant interaction (Fig. S2) and refer to it as sensitivity of the *FBA1* sequence to cleavage by the WHO endonuclease, whereas Δ*k* < 0.15 indicates resistance to cleavage.

**Figure 4.**
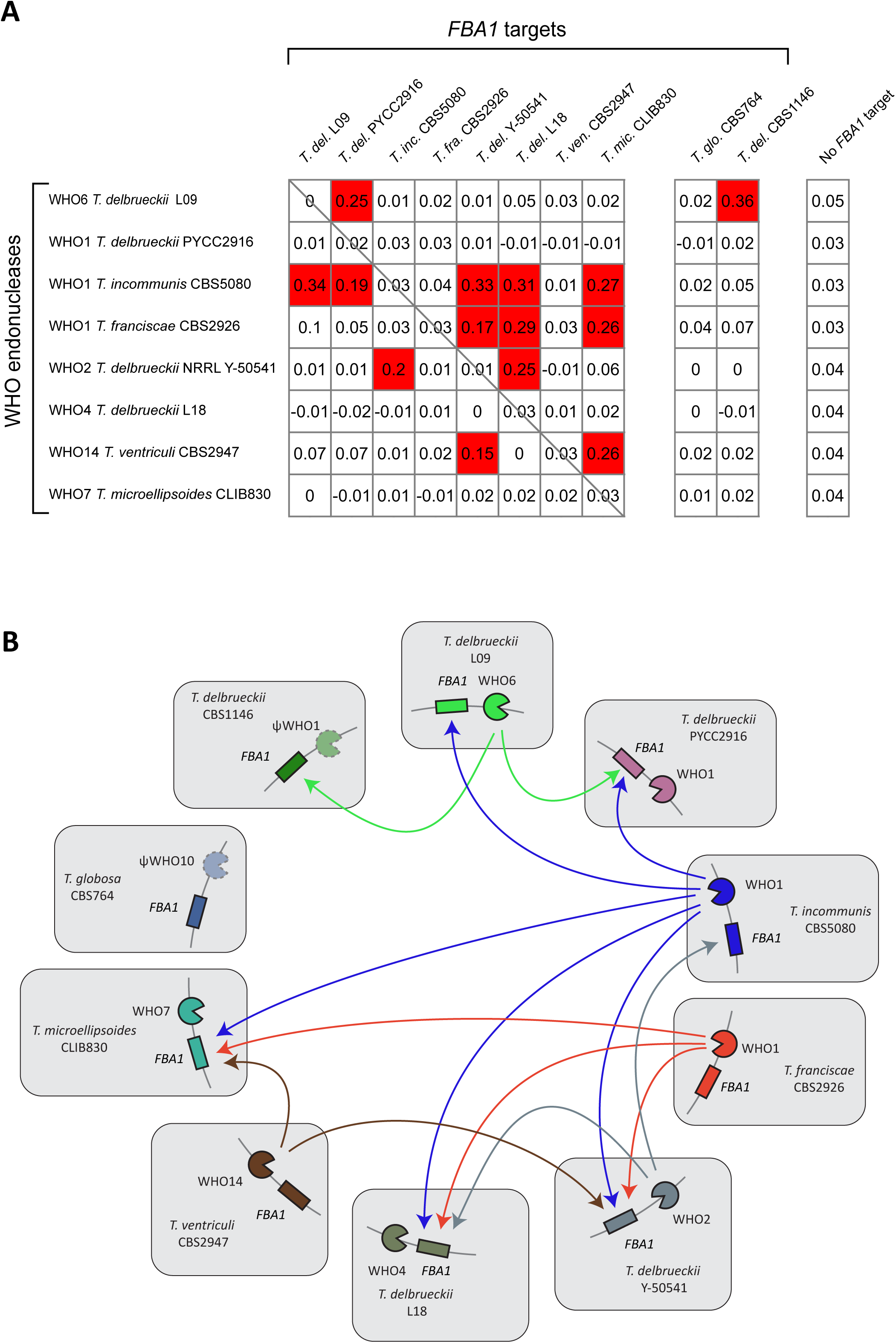
Allele-specific cleavage of *FBA1* targets by WHO endonucleases. **(A)** Matrix of interactions. Each WHO endonuclease (rows) was tested against each *FBA1* target sequence (columns). Cells show the difference (Δ*k*) between each strain’s growth rate in YPD (uninduced) and in YPD + β-estradiol (induced). Higher values of Δ*k* indicate greater sensitivity, and interactions scored as sensitive (Δ*k* ≥ 0.15) are highlighted in red. The diagonal line indicates interactions between *WHO* and *FBA1* genes that come from the same isolate of *Torulaspora* and are located directly beside each other in its genome. Strains CBS764 and CBS1146 did not have testable *WHO* genes. The rightmost column is a negative control showing Δ*k* values for strains that contain each *WHO* gene but do not have any target sequence inserted at *ho*. **(B)** Network representation of the interactions. Each gray box represents a complete WHO element, showing its ability to cleave, or to be cleaved by, other WHO elements. Arrows indicate sensitivity of the indicated *FBA1* sequence to the indicated WHO endonuclease (Δ*k* ≥ 0.15).

The matrix shows that there is extensive variation in the profiles of sensitivity and resistance to cleavage. Each *FBA1* sequence is resistant to the *WHO* gene that occurs directly downstream of it in the same genome (Fig. 4A, diagonal line), consistent with our hypothesis for the structure of WHO elements and their need to avoid self-cleavage. Three of the WHO endonucleases did not cleave any of the *FBA1* targets tested, whereas others were able to cleave between 2 and 5 of the 10 targets. WHO1 from *T. incommunis* CBS5080 was the most active endonuclease. It could cleave five targets, but notably it was unable to cleave two targets that were cleaved by other WHO endonucleases. Among the three WHO1 family members tested, two were very active but the third was inactive against all the targets tested. Among the *FBA1* targets, three were resistant to all the WHO endonucleases tested, whereas others were cleaved by 1–3 endonucleases. Three *FBA1* targets were sensitive to only one of the WHO endonucleases, but it was a different endonuclease in each case (Fig. 4A).

The matrix of interactions is drawn as a network in Figure 4B. Each gray box represents a complete WHO element present in one isolate of *Torulaspora*, consisting of a full-length *FBA1* gene and the adjacent *WHO* endonuclease gene. Arrows indicate the ability of each element to cleave, or to be cleaved by, other elements. The network indicates possible routes by which clusters of WHO elements could be formed. For example, the WHO element in *T. delbrueckii* CBS1146 is predicted to be vulnerable to invasion by the element from *T. delbrueckii* L09, which in turn is predicted to be vulnerable to invasion by the element from *T. incommunis* CBS5080 (Fig. 4B).

### Characterization of the WHO6 endonuclease recognition site in *FBA1*

LAGLIDADG endonucleases typically cleave DNA within a long (14–30 bp) and partially degenerate recognition sequence (5, 19). We characterized the sequence requirements for cleavage of the CBS1146 *FBA1* allele by WHO6 endonuclease, by first determining its approximate length and then systematically mutating every nucleotide position within this region.

To find the approximate length of the site, we inserted different-length regions from the sensitive *T. delbrueckii* CBS1146 *FBA1* allele into the *S. cerevisiae HO* locus by CRISPR-Cas9 and then assayed growth reduction after β-estradiol induction of WHO6 expression, on agar plates. The 83-bp region of *FBA1* used in the previous assays extends from positions −41 to +42, using a nucleotide numbering system in which −1/+1 marks the center of the cleavage site. We found that shorter regions of −31/+32, −21/+22 and -11/+22 bp were still cleaved, but that cleavage was lost when regions of −15/+15 and −21/+11 bp were tested (Fig. 5A). These results indicated that the minimal region is within the range of nucleotides −11/+22.

**Figure 5.**
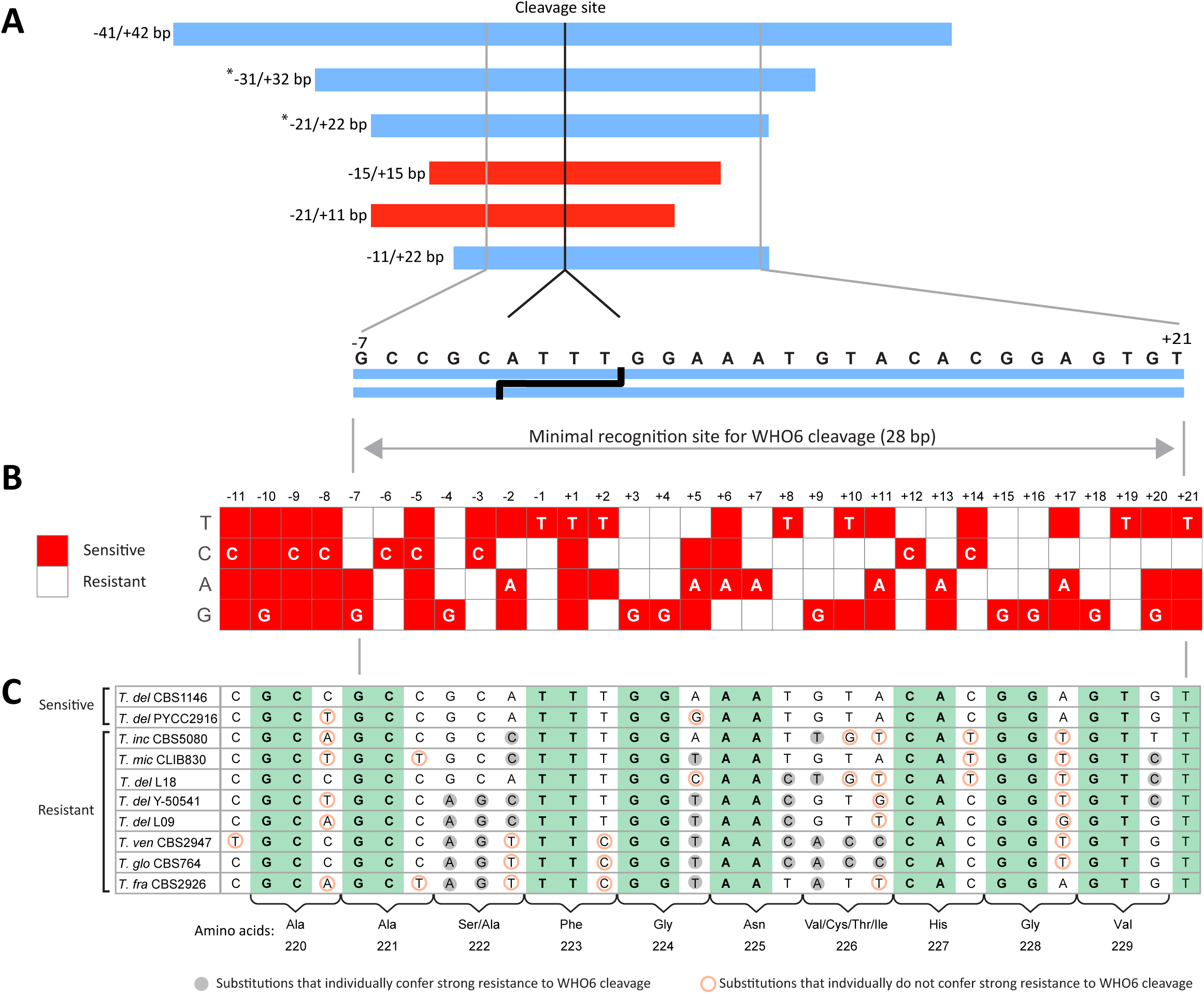
Characterization of the WHO6 recognition site in *FBA1*. (A) Delimitation of the *FBA1* region required for cleavage by WHO6. Bars represent fragments of the *T. delbrueckii* CBS1146 *FBA1* gene that were inserted into the *ho* locus of *S. cerevisiae*. Fragments that were cleaved after WHO6 induction are shown in blue, and fragments that were not cleaved are shown in red. The numbers of basepairs to the left (negative) and right (positive) of the center of the cleavage site are shown; the cleavage site was previously inferred to be at positions 666-669 relative to the start codon of *FBA1*, and to make a 4-bp overhang by analogy to HO and VDE endonucleases (9). Asterisks indicate regions that were tested using a WHO6 endonuclease with a FLAG epitope tag at its C-terminus; the other regions were tested using a WHO6 endonuclease without a FLAG tag. (B) Systematic mutagenesis of the minimal region. Red cells in the matrix indicate target sequences that led to significantly reduced growth (Δ*k* ≥ 0.15) after β-estradiol induction of *WHO6*, for all possible target sequences that differ from the CBS1146 *FBA1* sequence by 1 nucleotide in the region between positions -11 and +21. Bases introduced are indicated on the left. The original CBS1146 sequence is shown by white lettering. Δ*k* values are given in Table S1. (C) Comparison of *FBA1* allele nucleotide sequences in the minimal region. Amino acid translations are shown at the bottom. Green highlighting indicates nucleotides that are completely conserved. Circled nucleotides compare the allele sequences to the systematic mutagenesis results: gray circles mark nucleotides that conferred resistance to WHO6 cleavage, and red circles mark nucleotides that did not confer resistance, when introduced individually into the CBS1146 sequence.

To further delimit the region and its sequence specificity for cleavage, we made every possible single nucleotide change in the CBS1146 *FBA1* sequence between positions −11 and +21, and assayed their cleavage by using liquid growth assays. The results from this mutagenesis experiment (Fig. 5B) indicate that the minimal region required for cleavage of *FBA1* by WHO6 extends from nucleotides −7 to +21. The recognition sequence is 28 bp long and asymmetric around the cleavage site, as is typical of endonucleases with two LAGLIDADG motifs (5). WHO6 does not tolerate much variation in its target sequence. In the minimal region, 58 of the 84 mutations (69%) resulted in resistance (Fig. 5B; Table S1).

We compared the results from the mutagenesis experiment to our data (Fig. 4) on the cleavability of natural *Torulaspora FBA1* alleles by WHO6. The mutagenesis results can explain the resistance of each of the eight natural alleles that are resistant to WHO6: each of them has at least one nucleotide difference that confers resistance when introduced individually into the CBS1146 sequence (Fig. 5C, nucleotides highlighted by gray circles). The only other natural allele that is sensitive to WHO6, from *T. delbrueckii* PYCC2916, is identical to the CBS1146 sequence in the minimal region except for a change at position +5, which retains sensitivity when introduced individually into the CBS1146 sequence (Fig. 5B,C).

Most of the sequence variation among the natural isolates in the target region occurs at the third position of codons in the *FBA1* gene. Eight of the ten amino acids in the region are completely conserved among the isolates and are invariant at codon positions 1 and 2 (Fig. 5C, green highlighting), whereas amino acid positions 222 and 226 are variable. Many of the resistance-conferring substitutions between alleles are in codons 222 and 226, whereas many of the substitutions that do not remove sensitivity are at codon third positions. Among the 84 single-nucleotide mutations made in the minimal region (Fig. 5B), there are 60 that alter the *FBA1* amino acid sequence and these have an average Δ*k* of 0.068 (s.d. 0.071). The other 24 mutations are synonymous and their average Δ*k* is almost threefold higher at 0.189 (s.d. 0.076; *P* = 3 x 10^-8^ by T-test), so WHO6 can be said to tolerate more synonymous than nonsynonymous variation in its recognition sequence. Nevertheless, 8 of the 24 synonymous mutations resulted in resistance.

### Comparison to the HO and VDE recognition sites

The sequences of the minimal recognition sites for WHO6, HO (20) and VDE (21) endonucleases in their targets are compared in Figure 6. The lengths of the sites are 24 bp (HO), 28 bp (WHO6), and 31 bp (VDE). Each recognition site is asymmetric around its cleavage site. Even though these three endonucleases are closely related to each other, relative to the whole superfamily of LAGLIDADG endonucleases (6, 9), there is only low sequence similarity among their recognition sites. Only five nucleotides are conserved among all three endonuclease targets (positions −5, −4, +3, +5 and +7) in this alignment, which was centered by aligning the cleavage sites. The number of positions where a particular nucleotide is absolutely required for WHO6 cleavage is 13, which is higher than for HO and VDE (8 and 9 respectively; Fig. 6), so the WHO6 site is less degenerate.

**Figure 6.**
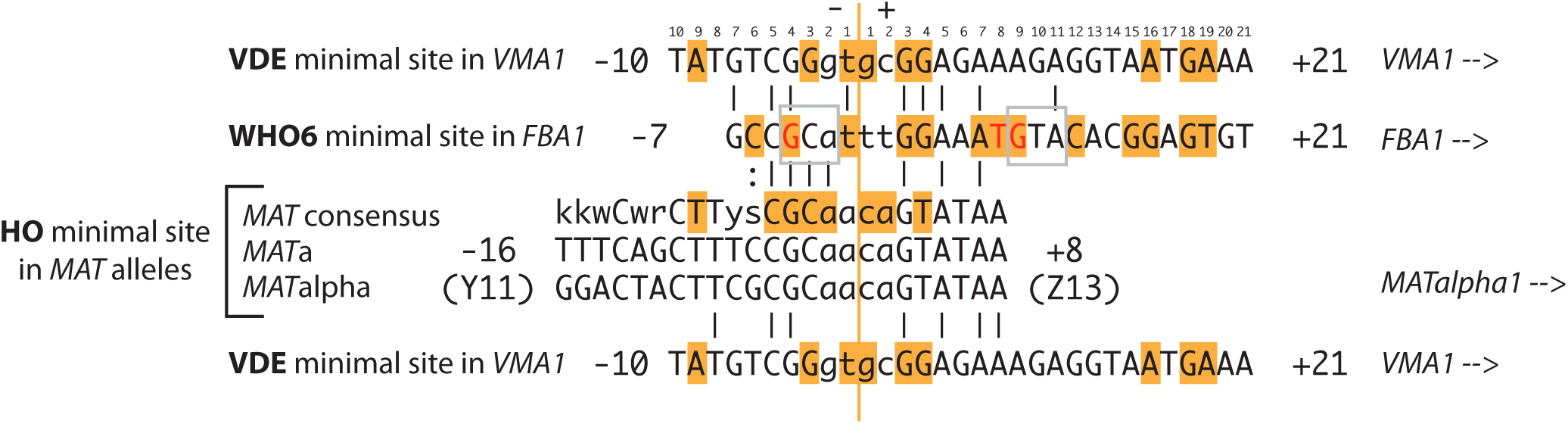
Comparison of the minimal recognition sites for WHO6, HO and VDE in their targets. Both *MAT*a and *MAT*α sites are shown for HO, and the VDE site is shown twice to facilitate comparison. Black vertical bars indicate matching bases. Sequences were aligned without gaps and are centered on the cleavage sites, which are known for HO and VDE and inferred for WHO6 (9). The orange line marks the center of the cleavage site, with the 4-bp overhangs shown by lowercase lettering (positions -2 to +2). Orange highlights indicate bases designated as critical for VDE cleavage by Gimble and Wang (21), for HO cleavage by Nickoloff *et al*. (20), and for WHO6 cleavage from our mutagenesis experiment (positions with Δ*k* < 0.15 for all 3 nucleotide substitutions). Three positions that are critical for WHO6 cleavage of the CBS1146 *FBA1* allele, but which vary in the sequences of other *FBA1* alleles that we found to be cleavable by other WHO endonucleases, are indicated by red font. The highly variable codons at amino acid positions 222 (GCA) and 226 (GTA) of *FBA1* are boxed. The sequence shown is the sense strand of each target gene. This alignment of the VDE and HO sites differs from the one presented by Gimble and Wang (21), who reversed the orientation of the HO site, introduced a gap, and did not align the cleavage sites.

### Positive selection on the WHO endonuclease cleavage site in *FBA1*

Interestingly, there are 3 positions in the CBS1146 *FBA1* allele where a particular nucleotide is absolutely required for cleavage by WHO6, but which are different in other *FBA1* alleles that we found could be cleaved by other WHO endonucleases (Fig. 4; Fig. 5C). Therefore, these 3 positions (−4, +8 and +9; red font in Fig. 6) are specifically required for cleavage by WHO6 but not by some other WHO endonucleases. These positions, which are determinants of the allele-specificity of cleavage, are located in the variable codons 222 and 226, or directly beside them. The high variability of these codons may be the result of natural selection on *FBA1* to avoid cleavage by particular WHO endonucleases.

If natural selection is acting on *FBA1* sequences to avoid cleavage by particular WHO endonucleases, it could result in positive selection on the sites in *FBA1* that determine sensitivity and resistance to cleavage. To investigate this possibility, we analyzed a larger set of 98 *Torulaspora* (mostly *T. delbrueckii*) genome sequences reported by Silva *et al*. (22), in addition to the genomes studied above. This dataset contained 50 different nucleotide sequences of the *FBA1* gene. We tested individual codon sites for evidence of positive selection (23, 24). At four codon sites, the nonsynonymous-to-synonymous substitution ratio (ω) is estimated to exceed 1, with high confidence (posterior probability > 0.95) (Fig. 7; Table S2). One of the codons showing the most significant evidence of positive selection is codon 226, which is the most variable codon within the recognition site (Fig. 5C) and contains a determinant of allele-specific cleavage (G at +9; Fig. 6). Another positively selected site is codon 232, which is just 3 codons downstream of the minimal recognition site. This result is consistent with the hypothesis that the *FBA1* alleles are under selection to avoid being cleaved.

**Figure 7.**
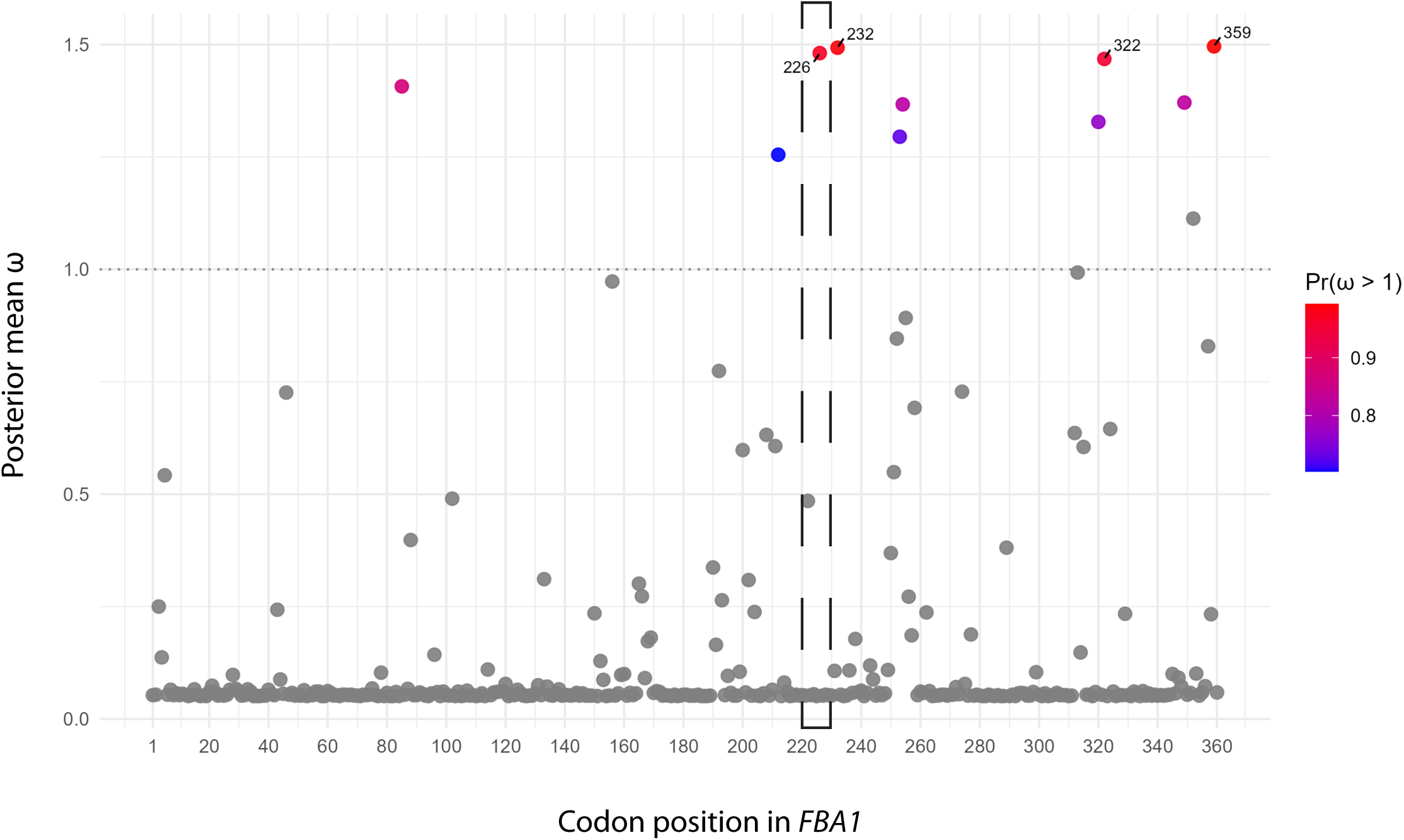
Positively selected sites in *Torulaspora FBA1* genes. Each point in the scatterplot is a codon in *FBA1*. The X-axis indicates codon positions along the *FBA1* gene, and the dashed rectangle shows the minimal WHO6 target site. The Y-axis indicates the best estimate (posterior mean) of the ratio of nonsynonymous to synonymous substitutions (ω) in each codon. The color scale indicates codons with a high posterior probability of having ω > 1, identified by using Bayes Empirical Bayes (BEB) inference (23). The four codons at which this probability exceeds 0.95 (Table S2) are labeled to indicate their positions.

## Discussion

Homing genetic elements tend to use highly conserved genes as their host genes, presumably a strategy that increases their cross-species invasiveness (1, 25, 26). For WHO elements the host gene is the aldolase gene *FBA1*, which is an essential, highly expressed yeast gene (27, 28). Adolase has a central role in metabolism because it catalyzes a step in the glycolysis pathway (cleavage of fructose-1,6-bisphosphate to glyceraldehyde 3-phosphate and dihydroxyacetone phosphate), and the corresponding reverse step during gluconeogenesis. Yeast aldolases are Class II aldolases, which use zinc ions for catalysis. The recognition site for WHO6 includes three residues (positions 226-228) that form part of the enzyme’s active site, by inference from the structure of the *Escherichia coli* enzyme ((29, 30); NCBI Conserved Domain database entry cd00946). His-227 binds the catalytic zinc ion and is conserved. Surprisingly, the neighboring position 226 in the active site is variable in *Torulaspora*, contains four different amino acids, and has undergone positive selection. Valine is the preferred amino acid at this position because it occurs in almost 99% of the Class II aldolase sequences in the NCBI database, whereas the other amino acids seen in *Torulaspora* are much rarer (Cys 1.04%; Thr 0.26%; Ile zero). This observation suggests that some alleles of *Torulaspora FBA1* have traded enzyme optimality for resistance to cleavage.

Our results confirm the hypotheses we previously proposed for the structure of WHO homing genetic elements and their interactions with *FBA1* (9, 11). We found that each *WHO* gene was unable to cause cleavage of the *FBA1* gene that occurs naturally directly upstream of it, which is consistent with the expectation that each WHO element will be resistant to self-cleavage. Most of the WHO endonucleases tested were able to cleave at least some *FBA1* alleles, and most of the *FBA1* alleles were cleaved by at least one WHO endonuclease. The apparent resistance of some *FBA1* alleles to all WHO endonucleases, and the failure of some WHO endonucleases to cleave any *FBA1* alleles, may be a consequence of the relatively low number of genes and targets tested. We know that *Torulaspora* species contain many more *FBA1* alleles and WHO variants than the ones we studied, and if we had assayed a larger dataset we might have detected cleavage involving the sequences that were inactive in our assays.

In interactions between different WHO elements, resistance is more common than sensitivity. Among the 72 *WHO*/*FBA1* combinations from different isolates of *Torulaspora*, only 14 (19%) showed cleavage (Fig. 4A). Even if we assume that the three WHO endonucleases that failed to cleave any targets failed for technical reasons, the cleavage rate is still only 33% (14 out of 42). Similarly, in our mutagenesis experiment, 69% of single-nucleotide changes in the 28-bp minimal recognition site resulted in resistance, and even 33% of synonymous changes resulted in resistance. Tolerance of synonymous substitutions by other LAGLIDADG endonucleases has been interpreted to be the result of selection to maximize their invasiveness (31), but this tolerance may be lower in WHO endonucleases than in other LAGLIDADG endonucleases because of selection on WHO elements to avoid self-cleavage.

We expect that any particular WHO element is under selection to have two properties (11). First, to maximize the element’s ability to spread through the population, its WHO endonuclease is under selection to be able to cleave the *FBA1* genes of strains that do not contain that element, i.e. strains that contain other WHO elements, or no element. The element’s endonuclease’s specificity should therefore be broad, but not so broad that it self-cleaves. Second, to minimize the element’s loss from the population by being a victim of homing, its *FBA1* is under selection to resist cleavage by all WHO endonucleases, while still coding for a functional aldolase enzyme.

This combination of selective pressures on each WHO element – to home, but to avoid being homed into – creates an arms race between *WHO* genes and their target site in *FBA1*. The signal of positive selection we detected at *FBA1* codon 226 is consistent with this evolutionary dynamic. So too is the finding that some nucleotides in the CBS1146 allele of the recognition site, including one in codon 226, are essential for cleavage by WHO6 but not by other WHO endonucleases. The DNA binding specificity of WHO endonucleases has probably co-evolved with their *FBA1* targets. We tested for evidence of positive selection on *Torulaspora WHO* genes but did not find any. However, our power to detect selection was probably low because of high sequence divergence among the different *WHO* families. The ability of WHO endonucleases to change their recognition sequence may have contributed to the evolutionary origin of HO endonuclease, which cleaves the mating-type locus, from a WHO-like ancestor (32).

WHO elements constitute a third category of homing genetic element, distinct from inteins and homing introns and considerably more rare. All three types of element use the same molecular mechanism of homing, but their genetic organization is different (11). Uniquely, when WHO elements integrate, they restore their host gene by replacing its 3’ half with an allelic variant of it, as opposed to disrupting the host gene and requiring it to be spliced. This difference in the element’s genetic organization leaves the restored host gene vulnerable to subsequent invasion by another WHO element with a different cleavage specificity, which has led to the emergence of an arms race and the formation of tandem clusters of WHO elements at the *FBA1* locus.

## Methods

### Yeast strains

Strain construction steps are summarized in Fig. 3. All *WHO*-expressing strains were constructed in *S. cerevisiae* strain MOY007 (Table S3; (33)). MOY007 is a derivative of the laboratory strain BY4742, made by integrating the plasmid FRP880 (Addgene #58437), which contains the synthetic β-estradiol-responsive transcription factor gene *LexA-ER-AD_B112* (16, 17), into the *his3*Δ1 locus of BY4742 by homologous recombination after *Pac*I linearization of the plasmid. *WHO* genes were synthesized by Twist Bioscience (South San Francisco, California, USA) and cloned into the integrating plasmid pRG634, which contains a β-estradiol inducible promoter (gift from Robert Gnügge; (17)) (Table S4). WHO-pRG634 vectors were linearized with *Asc*I and transformed into MOY007 using a lithium acetate transformation protocol (34) with selection on leucine dropout agar plus 100 μg/mL nourseothricin (HKI Jena). PCR with internal primers was used to check for the presence of the *WHO* gene, and pRG634 junctions were amplified to check for correct insertion into the *LEU2* locus (Table S5). Sequences (83 bp) from *Torulaspora FBA1* alleles were inserted into the *ho* locus using a two-plasmid CRISPR-Cas9 system with linear repair templates that had 150 bp of sequence identity to the *ho* locus on each side of the *FBA1* sequence (Table S6), and verified by sequencing. These steps generated haploid *S. cerevisiae* strains whose genomes contain both an inducible *WHO* gene and a potential target site from *Torulaspora FBA1* (Fig. 3D).

For delimitation and mutagenesis of the minimal target site, strain MOY195 (a derivative of MOY007 with *WHO6* integrated at the *LEU2* locus) was used. The WHO6-sensitive *FBA1* allele from *T. delbrueckii* CBS1146 was used to design 96 double stranded DNA repair templates in which each base between positions -11 and +21 of the target site was substituted by the three other possible bases (Fig. 5B; Table S1). For integration, the 96 repair templates each had 150 bp of sequence identity to the *ho* locus on each side of the 32-bp region from the *FBA1* target site (Table S5). They were synthesized by Twist Bioscience and inserted into the *ho* locus by CRISPR-Cas9 editing.

### WHO endonuclease sensitivity assays

For the assays of natural alleles shown in Fig. 4, at least two biological replicates were assayed, with two technical replicates of each, in liquid media. Detailed results are given in Table S7. For delimitation of the length of the minimal region (Fig. 5A) at least three biological replicates were assayed on solid media. For the mutagenesis experiment (Fig. 5B), 2-3 biological replicates were assayed, with two technical replicates each, in liquid media. For solid media assays, *S. cerevisiae* strains containing each *WHO* gene and *FBA1* target were streaked on YPD agar ± 2 μM β-estradiol (17), and photographed after 48 h incubation at 30° C. For liquid assays, overnight cultures were diluted to an OD600 of 0.1 in 200 μL of YPD or YPD + β-estradiol (1 μM) in 96-well plates, with duplicates for each biological replicate in each medium (no difference was seen between 2 μM and 1 μM β-estradiol inductions of a positive control). OD600 was measured by shaking for 2 minutes every 10 min for a 24 h period using a Synergy H1 microplate reader (BioTek). For each culture, the growth curve OD600 data was fitted to a logistic model using Rstudio (v2025.09.0+387), and the slope (*k*) of the growth curve at its inflection point, corresponding to the time when growth rate is maximal, was calculated. For each strain, technical replicates were averaged for each biological replicate, and the slope (*k*) was estimated independently for each biological replicate. The difference (Δ*k*) between the mean slopes of uninduced and induced cultures of each biological replicate was obtained by subtraction, and the reported Δ*k* value for the strain was calculated as the mean of Δ*k* among its biological replicates.

### Bioinformatic methods

For phylogenetic analysis, WHO protein sequences were aligned using Muscle and filtered with Gblocks in Seaview v5 (35). IQ-Tree v1.6.12 (36, 37) and ModelFinder (38) were then used for maximum-likelihood tree building. Tests for positive selection on *WHO* and *FBA1* genes were performed using site model analysis in CODEML (23, 24). Naive Empirical Bayes (NEB) and Bayes Empirical Bayes (BEB) inferences agreed on the 10 sites listed in Table S2. Amino acid frequencies at residue 226 of Class II aldolases in database sequences were calculated from the results of a BLASTP search against the NCBI nonredundant protein database, using *T. delbrueckii* aldolase as a query and retaining the top 4,978 hits from non-*Torulaspora* species; these sequences include aldolases from bacteria and diatoms as well as fungi.

## Acknowledgments

This work was supported by Research Ireland (20/FFP-A/8795). We thank Letal Salzberg, Padraic Heneghan, and Geraldine Butler for advice and comments, and Robert Gnügge for plasmids.

## Supplementary Information

**Figure S1.**
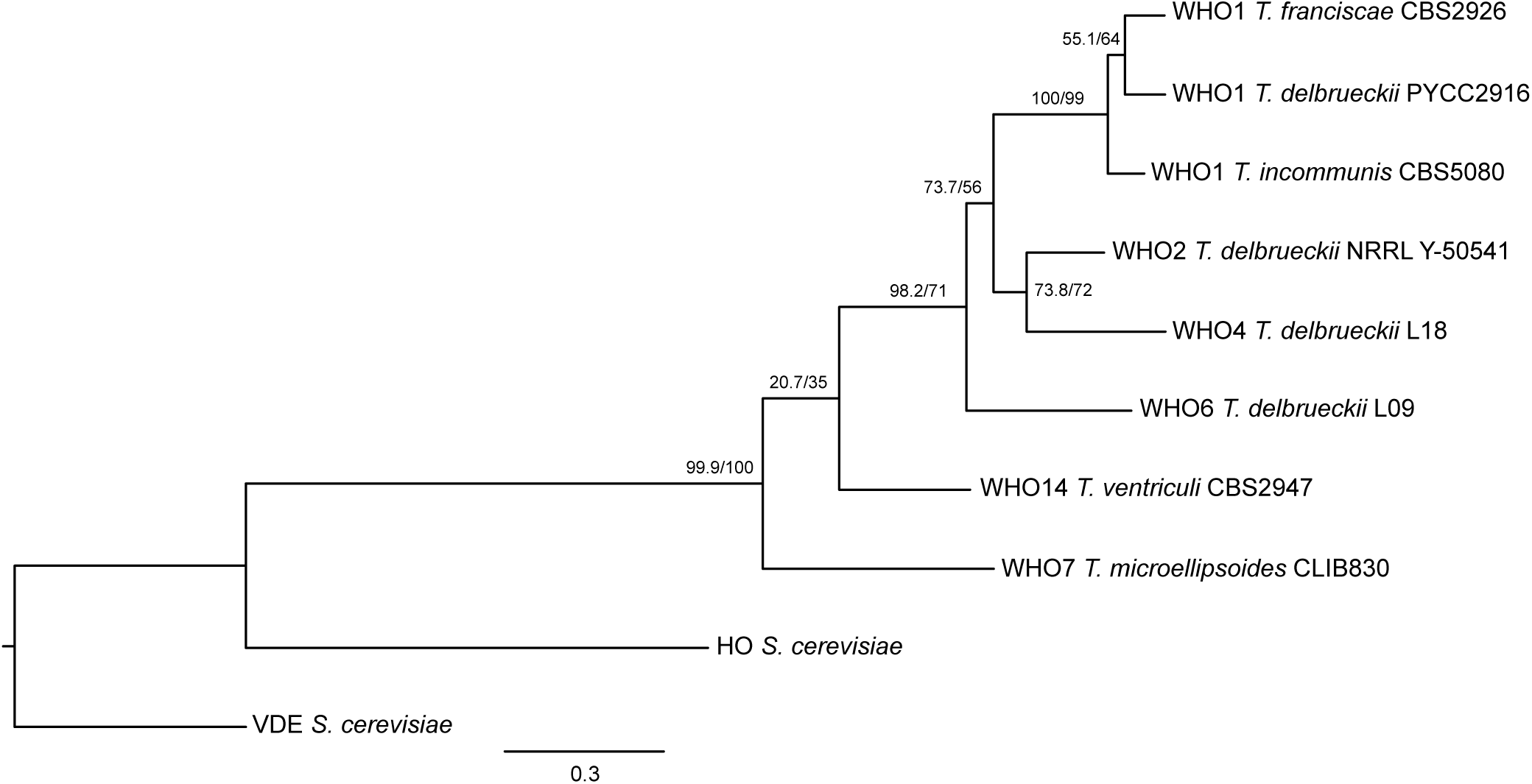
Phylogenetic relationship among the WHO proteins used in this study, HO and VDE. Protein sequences were aligned using Muscle and filtered with Gblocks in Seaview v5 (35). IQ-Tree v1.6.12 (36, 37) and ModelFinder (38) were then used for maximum-likelihood tree building. Numbers on branches show support values estimated by approximate Likelihood Ratio Test (aLRT; left) and 1000 ultrafast bootstraps (right). The tree was rooted using VDE.

**Figure S2.**
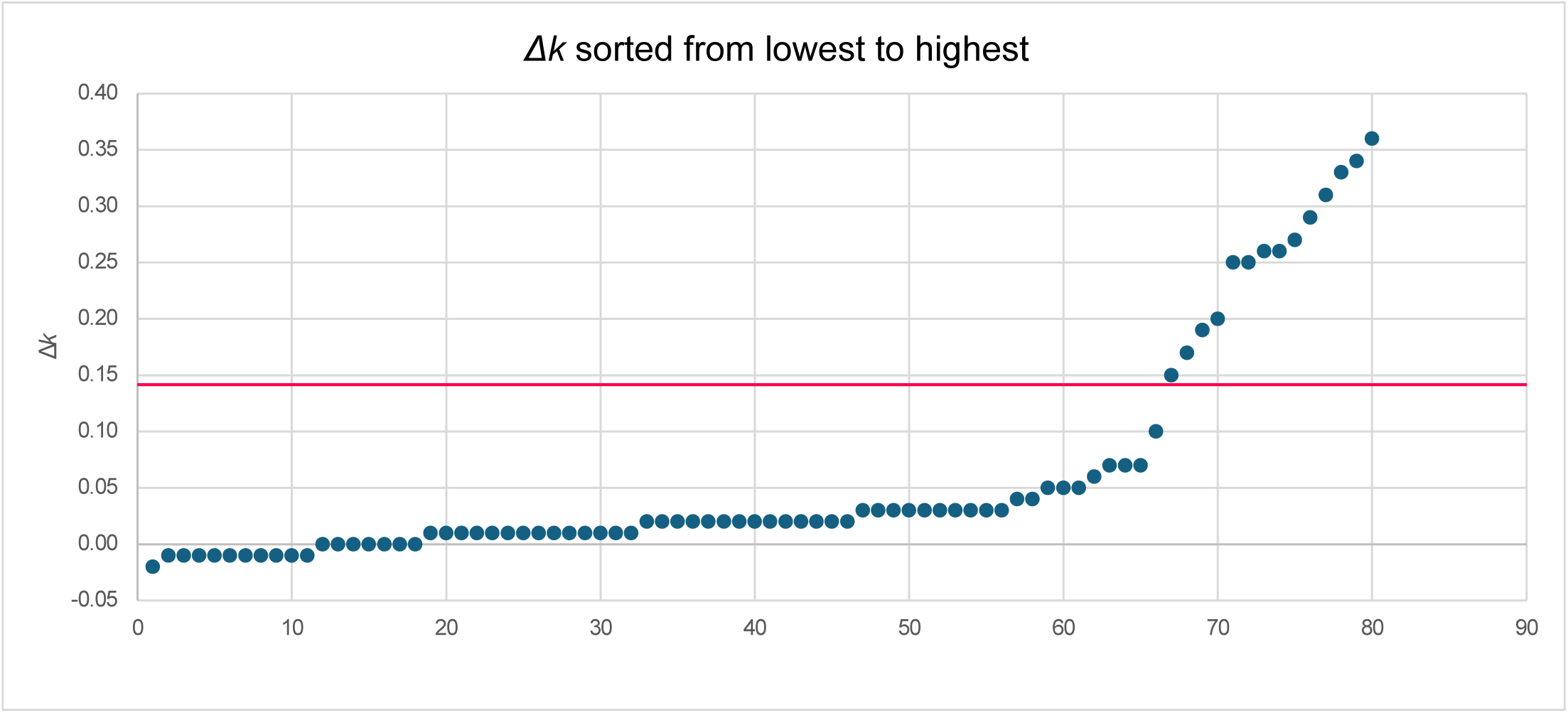
Growth rate differences (Δ*k* values) between mean uninduced and β-estradiol induced cultures, measured for the interactions between 8 WHO endonucleases and 10 *FBA1* alleles. The data is the same as in Figure 4A, with the 80 Δ*k* values sorted in increasing order on the X-axis. The red line indicates the threshold (Δ*k* ≥ 0.15) that was chosen to define sensitivity to cleavage.

**Supplementary Tables** are in a separate Excel file.

